# Craniofacial abnormalities from TNAP and PHOSPHO1 deletion highlight their dual role in bone and tooth mineralisation

**DOI:** 10.64898/2026.11.10.765640

**Authors:** Lucie E Bourne, Natalie Crump, Maya Karmali, Lucinda AE Evans, Aikta Sharma, Humayra Iqbal, Zain Ghani, Sonoko Narisawa, José Luis Millán, Brian L Foster, Colin Farquharson, Katherine A Staines

**Author notes:** **Corresponding author** Katherine Staines.

## Abstract

While tissue non-specific alkaline phosphatase (TNAP) and PHOSPHO1 are increasingly recognised as key cooperative regulators of postnatal skeletal mineralisation, the extent to which they interact during craniofacial morphogenesis has yet to be fully established. To elucidate these mechanisms, we generated mice harbouring a *Prx1*-mediated deletion of *Alpl* on a global *Phospho1*-null background, enabling a comprehensive assessment of phosphatase-mediated regulation of mineralisation across the developing craniofacial skeleton and dentoalveolar tissues. We show that whilst conditional deletion of *Alpl* had no effect on cranial and mandibular measurements, deletion of *Phospho1* resulted in a craniofacial hypoplasia characterised by frontonasal hypoplasia, reduced cranial vault height, and mandibular micrognathia, alongside increased intersphenoidal synchondrosis volume. This craniofacial hypoplasia phenotype was exacerbated by the dual deletion of *Alpl* and *Phospho1*, which resulted in further decreases in dentin volume and thickness. Together, these data indicate that TNAP and PHOSPHO1 play key roles in both craniofacial and dental development and add further evidence towards their synergistic role in supporting permissive biomineralisation.

## Introduction

The craniofacial complex encompasses the bones and soft tissues of the head and face, including the skull, jaws, teeth and facial muscles [1]. The skull, maxilla, and mandible develop through both endochondral and intramembranous ossification, whereas the transformation of the tooth bud into a fully mineralised tooth follow a distinct developmental process [1, 2]. Biomineralisation of bones and teeth is dependent on phosphatases that enable the formation and propagation of hydroxyapatite mineral within the collagenous matrix or enamel prisms, providing mechanical strength, rigidity and resistance to wear [3-5].

Tissue non-specific alkaline phosphatase (TNAP; encoded by *ALPL*) is expressed by osteoblasts [6], hypertrophic chondrocytes [7] and a variety of dentoalveolar cells – enamel-producing ameloblasts, dentin-producing odontoblasts, cementum-producing cementoblasts, and periodontal ligament (PDL) cells [8-11]. It is this ectoenzyme anchored to the outer membrane of matrix vesicles (MVs) that promotes matrix mineralisation by restricting the concentration of inorganic pyrophosphate (PPi) and dephosphorylating osteopontin, two potent inhibitors of mineralisation [12-14]. Pathogenic variants in the *ALPL* gene result in the extracellular accumulation of PPi within bone and teeth leading to hypophosphatasia (HPP), a rare inherited disease with a spectrum of clinical subtypes [15, 16]. Affected individuals exhibit skeletal defects including rickets, osteomalacia and fractures, whereas dental manifestations include acellular cementum deficiency, enamel alterations, thin and hypomineralised dentin, periodontal disease, and the premature loss of fully rooted deciduous and permanent teeth [8, 17-21]. Mineralised dental tissues are extremely sensitive to HPP with odontohypophosphatasia being the mildest subtype [22]. Whilst craniosynostosis is well established in the severe forms of the disease, such as infantile HPP, other craniofacial changes are poorly studied [23, 24].

PHOSPHO1, initially cloned from chick hypertrophic growth plate chondrocytes is now known to be expressed by osteoblasts, ameloblasts, odontoblasts and cementoblasts [4, 9, 25-29]. It resides within MVs and hydrolyses membrane-associated phosphoethanolamine and phosphocholine to initiate apatite crystal formation [27, 30, 31]. To our knowledge, no pathogenic, loss-of-function mutations in *PHOSPHO1* have been reported in humans although a patient with severe mineralisation defects had a heterozygous sequence variation c.95_97delCCT (p.Ser32del) in PHOSPHO1 and no detectable *ALPL* variants [32]. In osteoblasts and dental cells, the co-localisation of TNAP and PHOSPHO1 correspond with areas of mineralisation, suggesting that both phosphatases have complementary roles in the mineralisation process [9, 33].

Fundamental insights into the co-operativity of TNAP and PHOSPHO1 during the mineralisation process has advanced from cell culture observations [34] to the study of mice deficient in either phosphatase. Genetic ablation of *Alpl* in mice mimics infantile HPP, leading to severe abnormalities in skeletal and craniofacial mineralisation [35-37], as well as extensive dentoalveolar defects including the absence of acellular cementum and defective alveolar bone, enamel and root dentin mineralisation [11, 21, 38]. Since global *Alpl*-deficient mice die before weaning, *Alpl* conditional knockout mice instead have been generated to study late-onset HPP [39]. *Prx1-Cre* targeted deletion of *Alpl* in limb buds and craniofacial mesenchyme of adult mice resulted in hypomineralised alveolar bone, decreased root dentin thickness and an absence or reduction of acellular cementum [39]. More recently, we generated *Wnt1*^*Cre2*^; *Alpl*^*fl/fl*^ mice to conditionally delete *Alpl* in ectomesenchymal cells that make dentin, cementum, PDL and alveolar bone. This targeted approach, which also mimicked dental aspects of HPP, led to profound mineralisation defects in dentin, cementum, and alveolar bone [40].

In comparison to mice with a global or conditional deletion of *Alpl*, a deficiency in *Phospho1* leads to milder and less pronounced dentoalveolar abnormalities [41]. Specifically, *Phospho1* null mice present with enamel hypoplasia, thickened mantle dentin and reduced enamel, dentin and alveolar bone mineral densities [4, 9, 29, 41]. This reduction in dentin mineralisation was coincident with decreased MV mineralisation [9]. These abnormalities are similar to those observed in *Alpl* deficient mice; however, unlike *Alpl* null mice, the acellular cementum and PDL attachment remained intact in *Phospho1* knock-out mice [29, 41].

Despite growing evidence of their functional interplay in the mineralisation of the postnatal murine skeleton [42, 43], the combined roles of TNAP and PHOSPHO1 in the development of the craniofacial complex remain unclear. To address this, we generated a mouse model comprising a *Prx1*-driven deletion of *Alpl* on a global *Phospho1* background, permitting the evaluation of the critical roles of TNAP and PHOSPHO1 in controlling mineralisation during the development of the skull and dentoalveolar tissues.

## Results

### Deletion of *Phospho1* results in altered craniofacial measurements in 1-day-old mice

Mice with a *Prx1*-specific deletion of TNAP were bred onto a global *Phospho1*^-/-^ background (*Alpl*^*Prx1/Prx1*^*;Phospho1*^*-/-*^). Litters also consisted of wildtype (WT), *Prx1*-specific TNAP knockouts (*Alpl*^*Prx1/Prx1*^), *Phospho1* global knockouts (*Phospho1*^*-/-*^) and mice carrying a heterozygous *Prx1-*specific TNAP deletion on a *Phospho1*-null background (*Alpl*^*wt/Prx1*^*;Phospho1*^*-/-*^) [43].

Micro-computed tomography (µCT) analysis of skulls revealed that all postnatal day 1 (PN1) mice with a *Phospho1* deletion showed reduced bone surface area, in comparison to WT and *Alpl*^*Prx1/Prx1*^ mice (*p*<0.0001; Figure 1A). 3D-rendered µCT (Figure 1Bi) and alizarin red/alcian blue stained (Figure 1Bii) images demonstrate reductions in mineral content, which is most pronounced at the top of the skull. Anatomical landmark measurements (Suppl. Figure 1) revealed a reduced nasal bone (*p*<0.05; Figure 1Ci&ii) and frontal length (*p*<0.05; Figure 1Ciii) in these animals, which occurred independently of any change to total skull length (Figure 1Cix). Whilst skulls were also narrower (*p*<0.05; Figure 1Cxiii), a reduction in skull height was only observed in *Phospho1*^*-/-*^ animals (*p*<0.05; Figure 1Cx); however, all *Phospho1*^*-/-*^ containing genotypes had reduced mandibular parameters (*p*<0.05; Figure 1Cxi-xv). Despite this, no differences were observed in the intersphenoidal synchondrosis (ISS) or spheno-occipital synchondrosis (SOS) volume or thickness (Suppl. Figure 2A - D).

**Figure 1.**
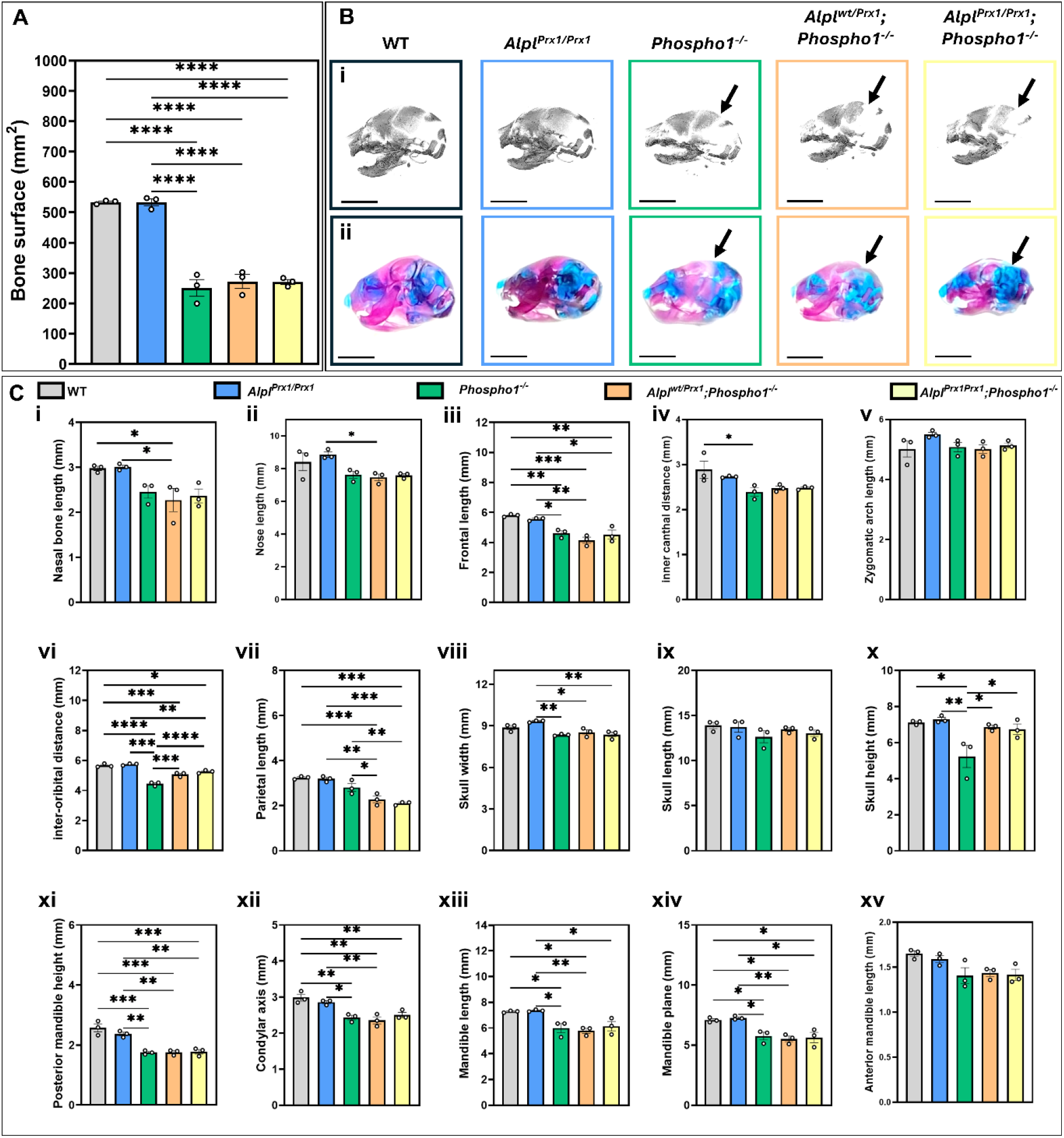
Altered craniofacial measurements in 1-day-old mice with Phospho1 deletion. (**A**) Total bone area of the skull in wild-type (WT), Alpl^Prx1/Prx1^, Phospho1^-/-^, Alpl^wt/Prx1^;Phospho1^-/-^ and Alpl^Prx1/Prx1^;Phospho1^-/-^ mice. (**B**) Representative (**i**) micro-computed tomography (μCT) renderings and (**ii**) whole mount images of the skull. Scale bar = 5 mm. Arrows indicate areas of reduced mineralisation. (**C**) Anatomical landmark measurements of the skull for (**i**) nasal bone length, (**ii**) nose length, (**iii**) frontal length, (**iv**) inner canthal distance, (**v**) zygomatic arch length, (**vi**) interorbital distance, (**vii**) parietal length, (**viii**) skull width, (**ix**) length, (**x**) height. Mandibular landmarks were also measured for (**xi**) posterior mandible height, (**xii**) condylar axis, (**xiii**) mandible length, (**xiv**) mandible plane, and (**xv**) anterior mandible length. Data presented as mean ± SEM with points showing individual animals (n=3/genotype). *= p<0.05, **=p<0.01, ***=p<0.001, ****= p<0.0001.

### Dual deletion of *Alpl* and *Phospho1* results in severe malformation of the skull in 3-week-old mice

By 3-weeks of age, all genotypes had reduced cranial bone surface area compared to WT, which progressively reduced in the single knockouts, to *Alpl*^*wt/Prx1*^*;Phospho1*^*-/-*^, and *Alpl*^*Prx1/Prx1*^*;Phospho1*^*-/-*^ (*p*<0.05; Figure 2A). Representative 3D-renderings of the skull reveal the reduced areas of mineralisation in *Phospho1*^-/-^, *Alpl*^*wt/Prx1*^*;Phospho1*^*-/-*^, and *Alpl*^*Prx1/Prx1*^*;Phospho1*^*-/-*^ mice, with complementary whole mount preparations highlighting the non-mineralised gap in the cranium occurring in *Alpl*^*Prx1/Prx1*^*;Phospho1*^*-/-*^ mice (Figure 2Bi&ii). Further, asymmetry in the nasal bones was noted in mice with a dual deletion of *Alpl* and *Phospho1* (Figure 2B, asterisk). Cranial and mandibular measurements were unaltered in *Alpl*^*Prx1/Prx1*^ mice, however *Phospho1*^*-/-*^-containing genotypes displayed a reduction in skull length-associated parameters (Figure 2Ci-vii, ix). Skull width and height were unchanged in all genotypes (Figure 2Cviii&x), resulting in a brachycephalic craniofacial skeletal phenotype, most severely observed with dual phosphatase deletion. Mandibular parameters were similarly reduced in *Phospho1*^*-/-*^-containing genotypes, with *Alpl*^*Prx1/Prx1*^*;Phospho1*^*-/-*^ mice displaying the shortest mandibles (Figure 2Cxi-xv).

**Figure 2.**
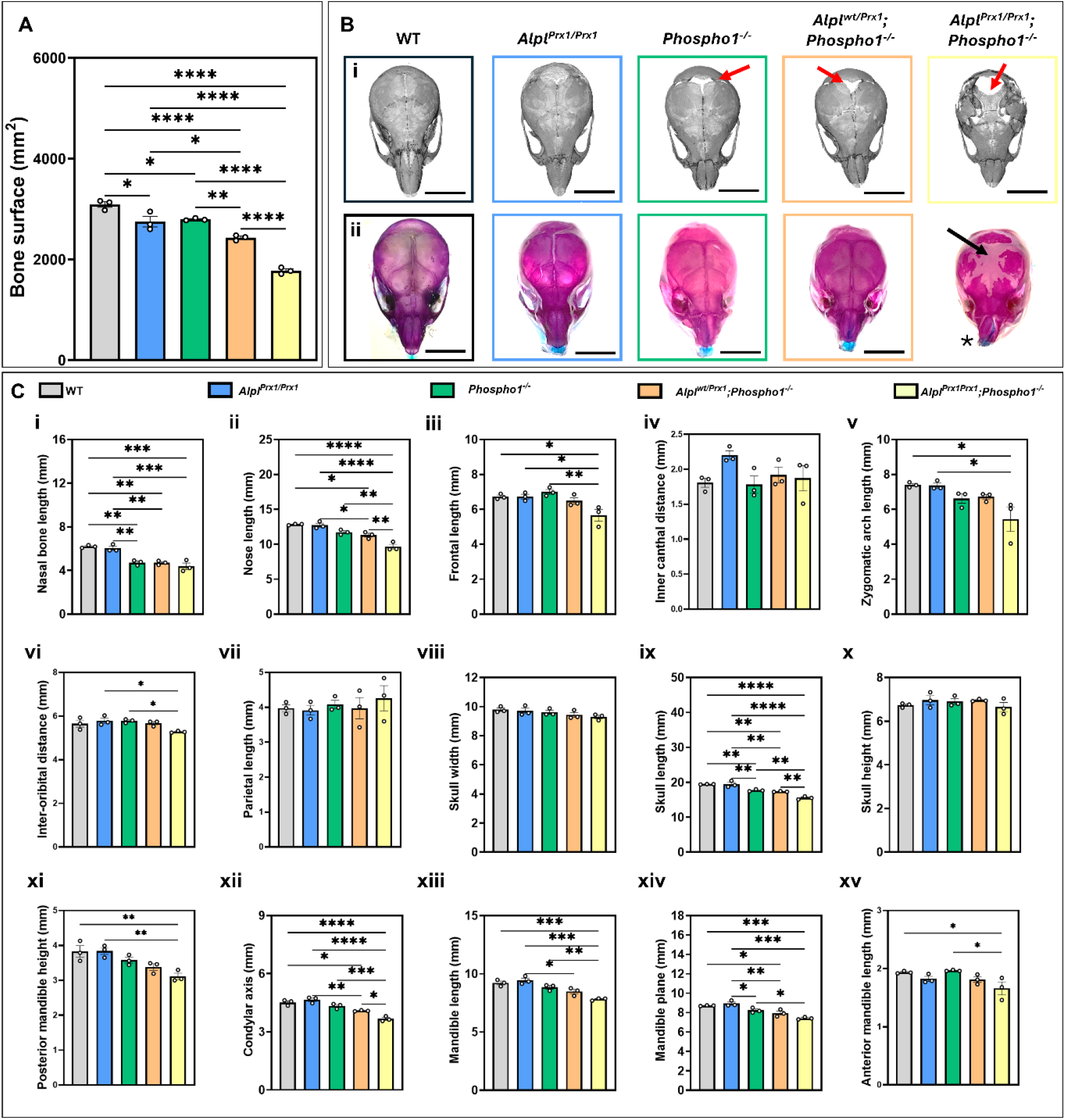
Skull malformation with dual Alpl and Phospho1 deletion at 3 weeks of age. (**A**) Total bone area of the skull in wild-type (WT), Alpl^Prx1/Prx1^, Phospho1^-/-^, Alpl^wt/Prx1^;Phospho1^-/-^ and Alpl^Prx1/Prx1^;Phospho1^-/-^ mice. (**B**) Representative (**i**) μCT-renderings and (**ii**) whole mount images of the skull. Scale bar = 5 mm. Arrows indicate areas of reduced mineralisation, asterisk indicates nasal bone asymmetry. (**C**) Anatomical landmark measurements of the skull for (**i**) nasal bone length, (**ii**) nose length, (**iii**) frontal length, (**iv**) inner canthal distance, (**v**) zygomatic arch length, (**vi**) inter-orbital distance, (**vii**) parietal length, (**viii**) skull width, (**ix**) length, (**x**) height. Mandibular landmarks were also measured for (**xi**) posterior mandible height, (**xii**) condylar axis, (**xiii**) mandible length, (**xiv**) mandible plane, and (**xv**) anterior mandible length. Data presented as mean ± SEM with points showing individual animals (n=3/genotype). *= p<0.05, **=p<0.01, ***=p<0.001, ****= p<0.0001.

At 3-weeks of age, differences in the synchondroses were restricted to the ISS, where deletion of *Phospho1* resulted in a significant increase in ISS volume in comparison to all other genotypes (Figure 3A & E, *p*<0.01). Moreover, the additional deletion of *Alpl* in *Alpl*^*Prx1/Prx1*^*;Phospho1*^*-/-*^ animals resulted in a significant reduction of the ISS volume in comparison to the WT, *Alpl*^*Prx1/Prx1*^, *Phospho1*^*-/-*^mice (Figure 3A & E, *p*<0.05) and in ISS thickness in comparison to *Phospho1*^*-/-*^mice (Figure 3B, *p*<0.05). Comparable changes, however, were not evident in the SOS (Figure 3C & D).

**Figure 3.**
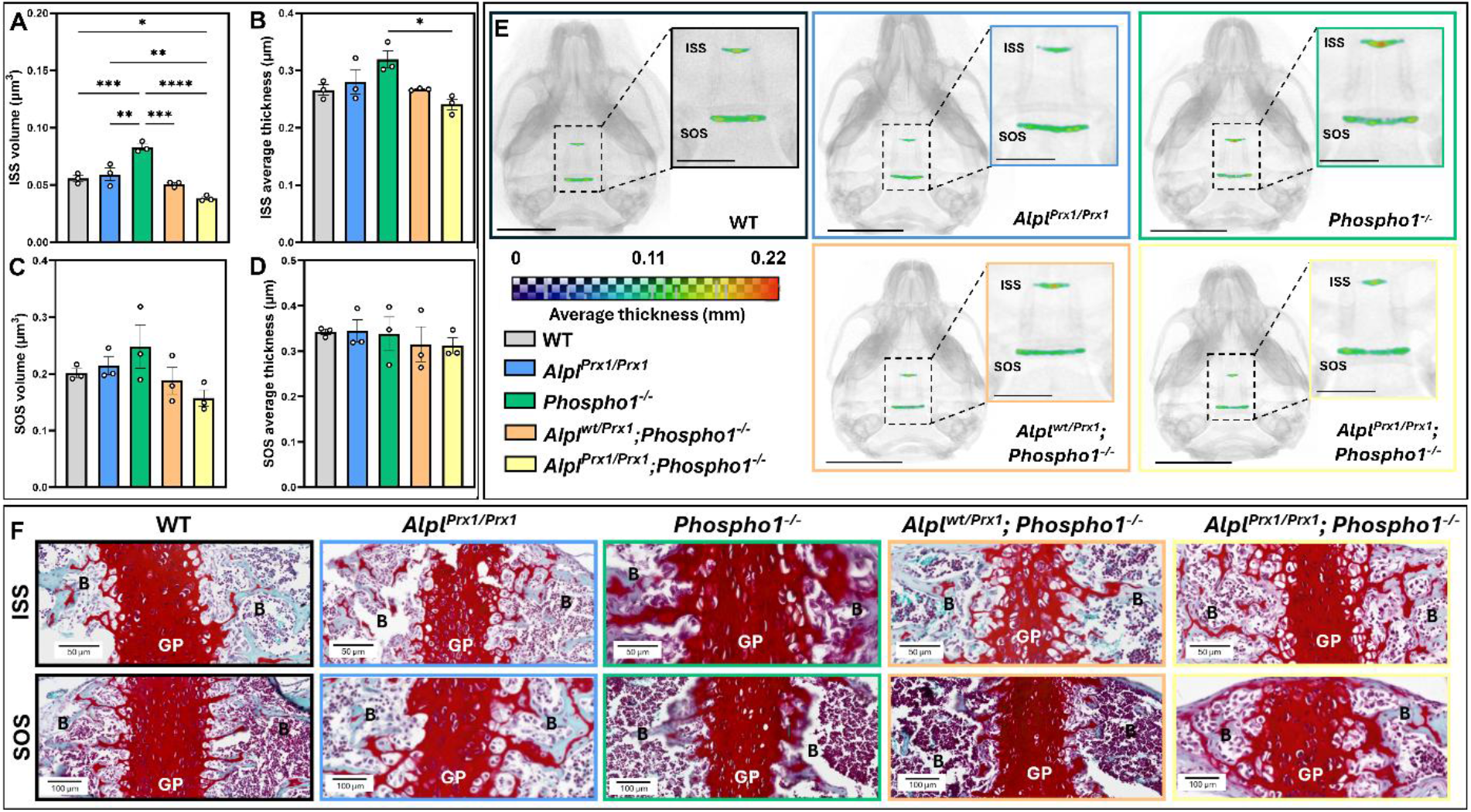
Aberrant intersphenoidal synchondrosis (ISS) formation with Phospho1 deletion and dual Alpl and Phospho1 deletion at 3 weeks of age. ISS (**A**) volume and (**B**) average thickness, and sphenooccipital synchondrosis (SOS) (**C**) volume and (**D**) average thickness in 3-week-old wild-type (WT), Alpl^Prx1/Prx1^, Phospho1^-/-^, Alpl^wt/Prx1^;Phospho1^-/-^ and Alpl^Prx1/Prx1^;Phospho1^-/-^ mice. (**E**) Representative 3D-rendered micro-computed tomography (μCT) images of the segmented ISS and SOS with average thickness heat map applied. Scale bar = 5 mm, or 2 mm in magnified images. (**F**) Representative Safranin O-stained histological sections of the ISS and SOS. GP = growth plate, B=bone. Scale bar = 50 µm (ISS) and 100 µm (SOS). Data presented as mean ± SEM with points showing individual animals (n=3/genotype). *= p<0.05, **=p<0.01, ***=p<0.001, ****= p<0.0001.

### Decreased dentin in the first molar of mice with dual deletion of *Alpl* and *Phospho1*

At 3-weeks of age, first mandibular molar crowns and roots are completely formed, and molars have erupted and are in function (Figure 4A & B). *Alpl*^*Prx1/Prx1*^*;Phospho1*^*-/-*^ mice had a reduced molar dentin volume (Figure 4C, *p*<0.05) and thickness (Figure 4D, *p*<0.05) in comparison to WT, *Alpl*^*Prx1/Prx1*^, *Phospho1*^*-/-*^ mice. *Alpl*^*wt/Prx1*^*;Phospho1*^*-/-*^ mice molar dentin trended towards decrease, suggesting a dose effect from increasing *Alpl* allele deletion. No significant differences in enamel volume or thickness were observed between genotypes (Figure 4E & F). Deletion of neither *Alpl* or *Phospho1* resulted in decreased alveolar bone volume or tissue mineral density (TMD) (Figure 4G & H, *p*<0.05). Incisor root analogue dentin on the lingual aspect was noticeably thinner in all genotypes lacking the *Phospho1* allele (Figure 4A & B).

**Figure 4.**
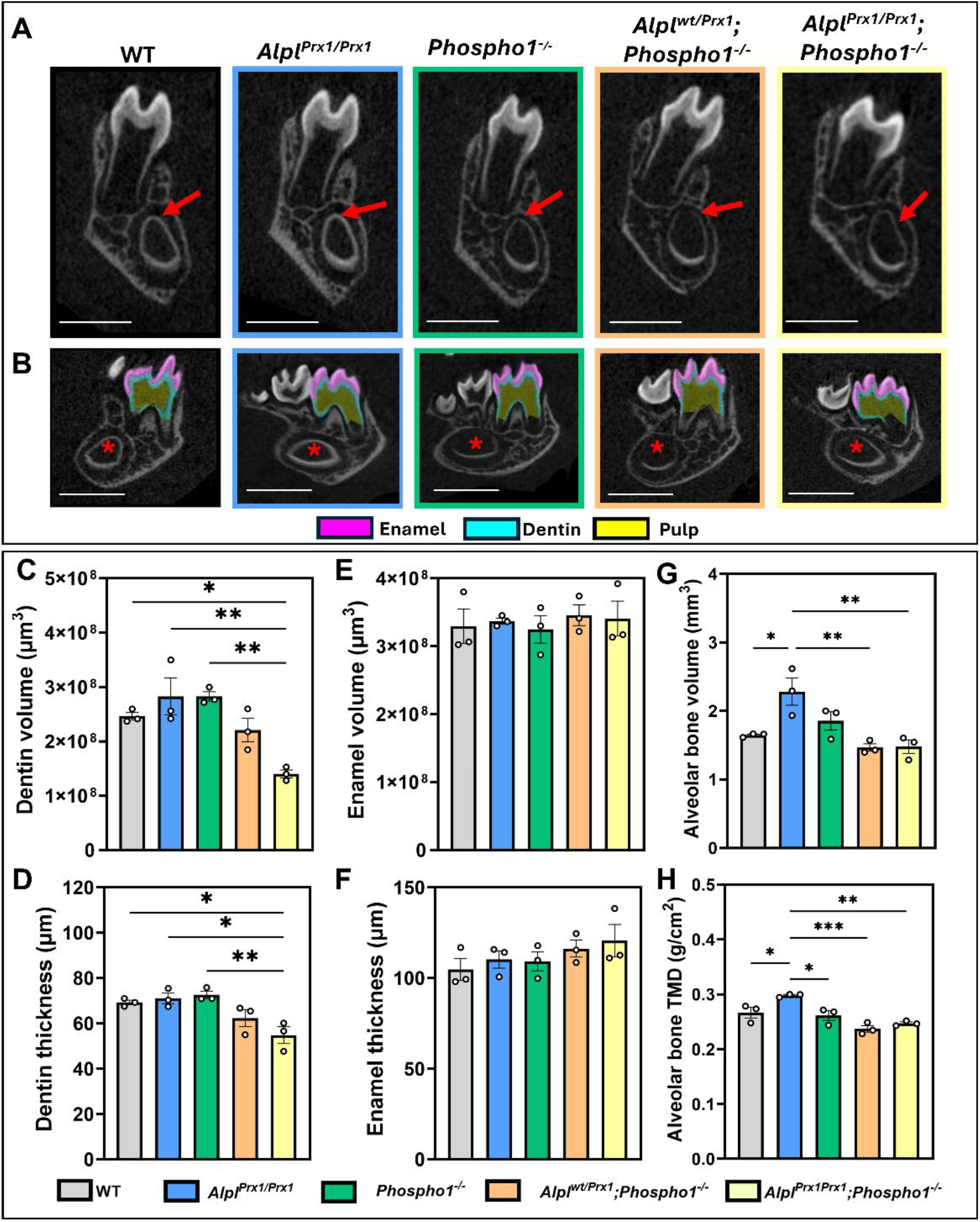
Dual deletion of Phospho1 and Alpl leads to reduced dentin at 3-weeks of age. (**A**) Representative μCT images of the M1 tooth and surrounding alveolar bone in wild-type (WT), Alpl^Prx1/Prx1^, Phospho1^-/-^, Alpl^wt/Prx1^;Phospho1^-/-^ and Alpl^Prx1/Prx1^;Phospho1^-/-^ mice at 3-weeks-of-age. Arrows indicate thinner dentin on the lingual part of the incisor in Phospho1-deficient animals. Scale bar = 1 mm. (**B**) Representative μCT images of the M1 segmentation of the enamel, dentin and pulp. Star indicates thinner dentin on the lingual part of the incisor in Phospho1-deficient animals. μCT analysis of dentin (**C**) volume and (**D**) average thickness, enamel (**E**) volume and (**F**) average thickness, and alveolar bone (**G**) volume and (**H**) mineral density (g/cm^2^). Data presented as mean ± SEM with points showing individual animals (n=3/genotype). *= p<0.05, **=p<0.01, ***=p<0.001, ****= p<0.0001.

### Minimal cranial and mandibular changes in 6-week-old mice of both sexes

Despite the changes described at earlier developmental time points, minimal cranial and mandibular changes were observed in 6-week-old mice (Suppl. Figure 3 & 4). Reductions in mandible length and plane persisted in *Phospho1*-containing genotypes, yet other anatomical measurements were no longer affected (Suppl. Figure 3 & 4). In the first molar, dentin volume and thickness differences present at 3 weeks were reduced or equalised by 6-weeks of age, whilst enamel volume and thickness were similarly not negatively affected by the loss of *Alpl* or *Phospho1* (Suppl. Figure 5).

## Discussion

Herein we utilised our murine model with dual deletion of *Alpl* and *Phospho1* to investigate the role of these two phosphatases in craniofacial development. We highlight that *Alpl* and *Phospho1* play key roles in supporting permissive biomineralisation in the skull. Loss-of-function mutations in *ALPL* cause HPP, a rare inherited disorder of mineralisation characterised by a broad spectrum of skeletal and dental abnormalities [15-17, 44]. Dental manifestations are particularly common and include defective dentin mineralisation, reduced cementum formation, and premature exfoliation of teeth [19, 21, 45-47]. Global deletion of *Alpl* in mice phenocopies severe infantile-onset HPP and results in profound craniofacial abnormalities, including skull bone hypomineralisation, altered cranial shape, and premature cranial suture fusion [11, 35, 36]. Global *Alpl* knockout mice also exhibit abnormal development of the cranial base synchondroses, with both the ISS and SOS displaying structural and developmental abnormalities [37]. Notably, *Alpl* deficiency appears to have a greater impact on cranial neural crest-derived bones than on bones of paraxial mesoderm origin, suggesting a tissue-selective requirement for TNAP during craniofacial development [37].

Due to the limitation of pre-weaning lethality with global *Alpl* deletion, several conditional *Alpl* knockout models have been developed and demonstrated similar craniofacial and mineralisation defects [39]. For example, neural crest-specific *Alpl* deletion (*P0-Cre*) results in abnormal development of the anterior (neural crest origin) but not posterior (mesoderm origin) cranial base, as well as an abnormal skull shape [48]. Further, *Col1a1-Cre* deletion of *Alpl* leads to craniofacial and postcranial skeletal defects, whilst *Wnt1-Cre* deletion of *Alpl* and caused craniofacial and dental defects [39, 40]. Collectively, these studies highlight a critical role for *Alpl* in regulating normal tooth, cranial bone, and cranial base development.

Despite this, our results herein show limited effects of *Alpl* deletion using a *Prx1-Cre*, which is expressed during early development in limb bud mesenchyme of the appendicular skeleton and in a subset of craniofacial/dental mesenchyme [49]. This is somewhat surprising given the clear skeletal phenotype we recently observed in the long bones of these mice [43], and given that our global deletion of *Phospho1* resulted in a craniofacial hypoplasia phenotype characterised by frontonasal hypoplasia, reduced cranial vault height, and mandibular micrognathia. These findings are consistent with the profound mineralisation defects previously reported in *Phospho1*-deficient mice, which exhibit marked impairment of skeletal mineralisation and bone formation [30]. However previous work examining the craniofacial and dental phenotypes using *Prx1-Cre* deletion of *Alpl* has been contradictory [39, 40, 50]. The reasons for this are unclear and may be underpinned by insufficient *Cre* recombination in the craniofacial mesenchyme, or insufficient *Prx1-Cre* expression in *Alpl* positive cells in these tissues [39]. Further it is possible that any lack of TNAP in the *Prx1* positive cells are compensated for by other mechanisms such as *Phospho1*. Our data showed that the dual deletion of *Alpl* and *Phospho1* exacerbated the craniofacial hypoplasia phenotype observed in *Phospho1*-deficient mice alone, resulting in further reductions in craniofacial dimensions. This therefore supports the theory that the absence of a significant craniofacial phenotype in our *Alpl*^*Prx1/Prx1*^ mice is likely due to *Prx1-Cre* not being expressed in a sufficient proportion of *Alpl*-expressing cells within the teeth, cranial base synchondroses, or temporomandibular joint.

Examination of the cranial base revealed a selective enlargement of the ISS in *Phospho1*-deficient mice, with no differences seen with *Prx1*-directed *Alpl* deletion. This is in contrast to the global *Alpl* knockout mouse in which an abnormal ISS was characterised by a widely expanded hypertrophic zone and further highlights the limitations of the *Prx1-Cre* deletion [37, 39]. However interestingly, in our double knockout animals, the ISS dimensions were significantly reduced despite the more severe craniofacial phenotype. These findings suggest that synchondrosis size does not necessarily directly reflect growth potential. Rather, ISS enlargement in *Phospho1*-deficient mice may represent delayed cartilage maturation, with accumulation of hypertrophic chondrocytes and prolonged growth plate-like activity, analogous to the enlarged hypertrophic zones previously reported in the appendicular skeleton of *Phospho1*-deficient mice [43]. In contrast, combined loss of PHOSPHO1 and TNAP may impair the integrity and maintenance of the synchondrosis growth centre itself, leading to depletion or disorganisation of the cartilage template, reduced synchondrosis size, and impaired cranial base elongation. Under this model, the enlarged ISS in *Phospho1* knockout mice may represent a partially compensatory mechanism that helps preserve cranial base growth despite defective mineralisation. Additional loss of *Alpl* abolishes this compensatory response, resulting in a smaller, dysfunctional ISS and a more severe craniofacial phenotype. Additional experiments are warranted to clarify effects on cranial base synchondroses and how these may be influencing craniofacial growth.

In dentoalveolar tissues, we observed limited differences with conditional *Alpl* or global *Phospho1* deletion, which contrasts with previous studies. In *Phospho1* knockout mice, histological analyses by Zweifler and colleagues identified an expanded hypomineralised region within the outer dentin, corresponding to the mantle dentin [29]. This has been further examined with µCT in which alterations in enamel volume, dentin volume and density, as well as reduced alveolar bone density were observed in *Phospho1*-deficient mice [9, 29]. The lack of differences observed in this present study may reflect the substantially younger age of the animals analysed and the developmental timing of mineralisation defects. It may further reflect the limitation of using µCT analyses which may not be able to detect subtle local changes in the dentoalveolar tissues, especially the cementum. The relatively normal molar dentition in the *Alpl*^*Prx1/Prx1*^ mice herein further adds to our evidence that *Prx1-Cre* deletion may not recapitulate the full spectrum of HPP phenotypes. This is further exemplified by our analyses of the alveolar bone in 3-week-old animals in which we saw a significant increase with *Alpl* deletion which is at odds with previous studies [41]. Work by McKee and colleagues examined mice lacking *Phospho1* and one allele of *Alpl* (*Phospho1*^*-/-*^*;Alpl*^*+/-*^) and found reduced dental mineralisation in the incisors by transmission electron microscopy [9]. This is consistent with our findings in the double *Alpl* and *Phospho1* knockout mice which exhibited reduced dentin thickness and volume, suggesting a profound disturbance in tooth mineralisation. Further, Mckee reported that in *Phospho1*^*-/-*^*;Alpl*^*+/-*^ mice, there was a complete lack of MVs in the mantle dentin which was postulated to underpin the mineralisation defect observed due to a lack of key enzymes involved in P_i_ and PP_i_ processing [9]. Further studies could however examine whether this is also the case in our double knockout animals.

In summary, our data indicate that dual deletion of TNAP and PHOSPHO1 leads to craniofacial dysmorphology and dental defects, supporting the model that these phosphatases function synergistically to support permissive biomineralisation.

## Methods

### Animals

All tissue isolation and experimental procedures were performed in accordance with the UK Animals (Scientific Procedures) Act of 1986 and regulations set by the UK Home Office and local institutional guidelines at the University of Brighton.

Tissue-specific *Alpl*^*Prx1/Prx1*^ mice were generated using the *Prx1-Cre* transgene as previously reported [39]. The generation and characterisation of Phospho1-R74X mutant mice (Phospho1^m1Jlm^, here referred to as *Phospho1*^*−/−*^) were also previously described [42]. To generate mice lacking global PHOSPHO1 and *Prx1*-conditional TNAP, *Phospho1−/−* and *Alpl*^*Prx1/Prx1*^ were crossed to generate double mutant (*Alpl*^*Prx1/Prx1*^;*Phospho1*^*−/−*^) and *Alpl*-heterozygous (*Alpl*^*wt/Prx1*^;*Phospho1*^*−/−*^) mice, as previously described [43]. Genotyping was conducted by Transnetyx using their automated genotyping service. Whole skulls were collected from both male and female mice at PN1, 3-weeks of age, following euthanasia by Schedule 1. Analyses were conducted blindly to minimise the effects of subjective bias. While *Alpl*^*Prx1/Prx1*^*;Phospho1*^*-/-*^ animals survived, they displayed severe post-weaning growth restriction and failed to thrive relative to littermate controls as previously reported [43]. Ethical endpoints mandated under UK animal welfare legislation precluded maintenance of these mice beyond 3 weeks of age. Accordingly, assessment of the craniofacial phenotype in skeletally mature mice at 6-weeks of age were limited to the other experimental genotypes.

### Whole mount staining

PN1 and 3-week-old animals were skinned, eviscerated and fixed in 95% ethanol overnight, as previously described [51]. Acetone (100%) was replaced for 1-2 days. Preparations were incubated in 0.03% alcian blue (w/v), 80% ethanol, 20% glacial acetic acid for 1-3 days, before destaining in two changes of 70% ethanol and incubating in 95% ethanol overnight. Samples were pre-cleared in 1% potassium hydroxide (KOH) solution overnight at 4°C and incubated in 0.005% alizarin red (in 1% KOH) at 4°C for 1-3 days. Specimens were cleared in 1% KOH and stored in 50% glycerol:50% KOH (1%) before imaging.

### µCT analysis

Scans of PN1 and 3-week-old (n=3/genotype), and 6-week-old (n=5-7/genotype/sex) skulls were performed using a Skyscan 1172 X-Ray microtomograph (Bruker). Tomographic scans (200 µA, 0.5 mm aluminium filter, 0.6° rotation angle) were acquired with a voxel size of 10 µm (40 kV) for PN1 animals; and 13.5 µm (50 kV) for 3-and 6-week-old animals (40 kV). The projection images were reconstructed using NRecon (Bruker, version 1.7.4.6), datasets orientated and aligned in DataViewer (Bruker, version 1.5.6.2), and regions of interest were segmented in CTAn (Skyscan).

Bone surface area of the entire skull was performed on binarised images using a minimum threshold of 80 and 3D measurements were obtained in CTAn. Anatomical measurements were performed in Avizo software (ThermoFisher Scientific), as previously described (Suppl. Figure 1) [37, 52]. To evaluate volumes (µm^3^) and average thicknesses (µm, parallel plate model) of (i) the ISS and SOS, and (ii) the dentin and enamel of the first molars (M1), these were manually segmented using Avizo software (ThermoFisher Scientific) [53]. To generate false-colour thickness heatmaps, segmentations were resampled, and the Avizo ‘thickness map’ module with the ‘physics’ Lookup Table applied.

### Histology

Skulls were fixed for 24 hours in 4% paraformaldehyde before being decalcified in 10% ethylenediaminetetraacetic acid for 21 days at 4oC, and wax-embedded. Sagittal sections (6 μm) were stained with Safranin O/Fast green using standard procedures.

### Statistical analysis

Analyses were performed using GraphPad Prism software (version 10.1.2). Normal distribution of data was assessed using the Shapiro-Wilk normality test. For comparison between genotypes, a one-way ANOVA followed by Tukey’s post hoc analysis was used to test for significance. Results are presented as the mean ± standard error of the mean (SEM).

## Data Availability

The datasets used and/or analysed during the current study are available from the corresponding author on reasonable request.

## Acknowledgements

The authors would like to thank the Bioresources Unit at the University of Brighton for the assistance with the care of animal models in this study. We also thank Prof. Denis Headon (Roslin Institute) for his discussions on the breeding strategy. We are also thankful to Dr Mark Hopkinson (Royal Veterinary College) for technical assistance. The authors would like to acknowledge the UKRI Medical Research Council (MRC) for funding to KAS (MR/V033506/1 & MR/R022240/2), the Biotechnology and Biological Sciences Research Council (BBSRC) for Institute Strategic Programme Grant Funding to CF (BBS/E/D/10002071 & BBS/E/RL/230001C), and NIDCR (R01DE032334) for funding BLF.

For the purpose of open access, the authors have applied a Creative Commons Attribution (CC-BY) license to any Author Accepted Manuscript version arising from this submission.

## Author Contributions

LEB, KAS, CF, JLM, SN, BLF designed research; LEB, NC, LAEE, MK, AS, HI, ZG performed research; LEB, NC, KAS analysed data; LEB, NC, CF, KAS wrote the manuscript. All authors reviewed and edited the manuscript.

## References

1. D’Souza RN, Ruest L-B, Hinton RJ, Svoboda KKH. Development of the Craniofacial Complex. In: Bronner F, Farach-Carson MC, Roach HI, editors. Bone and Development. London: Springer London; 2010. p. 153–81.

2. Novacescu D, Dumitru CS, Zara F, Raica M, Suciu CS, Barb AC, et al. The Morphogenesis, Pathogenesis, and Molecular Regulation of Human Tooth Development-A Histological Review. Int J Mol Sci. 2025;26(13). Epub 20250627. doi: 10.3390/ijms26136209. PubMed PMID: 40649989; PubMed Central PMCID: PMCPMC12250405.

3. Donnelly E, Chen DX, Boskey AL, Baker SP, van der Meulen MC. Contribution of mineral to bone structural behavior and tissue mechanical properties. Calcif Tissue Int. 2010;87(5):450–60. Epub 20100822. doi: 10.1007/s00223-010-9404-x. PubMed PMID: 20730582; PubMed Central PMCID: PMCPMC2965269.

4. Pandya M, Rosene L, Farquharson C, Millán JL, Diekwisch TGH. Intravesicular Phosphatase PHOSPHO1 Function in Enamel Mineralization and Prism Formation. Front Physiol. 2017;8:805. Epub 20171017. doi: 10.3389/fphys.2017.00805. PubMed PMID: 29089903; PubMed Central PMCID: PMCPMC5651051.

5. Millán JL. The role of phosphatases in the initiation of skeletal mineralization. Calcif Tissue Int. 2013;93(4):299–306. Epub 20121127. doi: 10.1007/s00223-012-9672-8. PubMed PMID: 23183786; PubMed Central PMCID: PMCPMC3594124.

6. Hasegawa T. Ultrastructure and biological function of matrix vesicles in bone mineralization. Histochem Cell Biol. 2018;149(4):289–304. Epub 20180206. doi: 10.1007/s00418-018-1646-0. PubMed PMID: 29411103.

7. Farquharson C, Hesketh JE, Loveridge N. The proto-oncogene c-myc is involved in cell differentiation as well as cell proliferation: studies on growth plate chondrocytes in situ. J Cell Physiol. 1992;152(1):135–44. doi: 10.1002/jcp.1041520118. PubMed PMID: 1618915.

8. Foster BL, Nagatomo KJ, Nociti Jr FH, Fong H, Dunn D, Tran AB, et al. Central role of pyrophosphate in acellular cementum formation. PLoS One. 2012;7(6):e38393.

9. McKee MD, Yadav MC, Foster BL, Somerman MJ, Farquharson C, Millan JL. Compounded PHOSPHO1/ALPL deficiencies reduce dentin mineralization. J Dent Res. 2013;92(8):721–7. Epub 20130521. doi: 10.1177/0022034513490958. PubMed PMID: 23694930; PubMed Central PMCID: PMCPMC3711567.

10. Hotton D, Mauro N, Lézot F, Forest N, Berdal A. Differential expression and activity of tissue-nonspecific alkaline phosphatase (TNAP) in rat odontogenic cells in vivo. J Histochem Cytochem. 1999;47(12):1541–52. doi: 10.1177/002215549904701206. PubMed PMID: 10567438.

11. Foster BL, Nagatomo KJ, Tso HW, Tran AB, Nociti FH, Jr., Narisawa S, et al. Tooth root dentin mineralization defects in a mouse model of hypophosphatasia. J Bone Miner Res. 2013;28(2):271–82. doi: 10.1002/jbmr.1767. PubMed PMID: 22991301; PubMed Central PMCID: PMCPMC3541444.

12. Addison WN, Azari F, Sørensen ES, Kaartinen MT, McKee MD. Pyrophosphate inhibits mineralization of osteoblast cultures by binding to mineral, up-regulating osteopontin, and inhibiting alkaline phosphatase activity. J Biol Chem. 2007;282(21):15872–83. Epub 20070323. doi: 10.1074/jbc.M701116200. PubMed PMID: 17383965.

13. Meyer JL. Can biological calcification occur in the presence of pyrophosphate? Arch Biochem Biophys. 1984;231(1):1–8. doi: 10.1016/0003-9861(84)90356-4. PubMed PMID: 6326671.

14. Harmey D, Johnson KA, Zelken J, Camacho NP, Hoylaerts MF, Noda M, et al. Elevated skeletal osteopontin levels contribute to the hypophosphatasia phenotype in Akp2(-/-) mice. J Bone Miner Res. 2006;21(9):1377–86. doi: 10.1359/jbmr.060619. PubMed PMID: 16939396.

15. Millán JL, Whyte MP. Alkaline Phosphatase and Hypophosphatasia. Calcif Tissue Int. 2016;98(4):398–416. Epub 20151121. doi: 10.1007/s00223-015-0079-1. PubMed PMID: 26590809; PubMed Central PMCID: PMCPMC4824800.

16. Whyte MP, Coburn SP, Ryan LM, Ericson KL, Zhang F. Hypophosphatasia: Biochemical hallmarks validate the expanded pediatric clinical nosology. Bone. 2018;110:96–106. Epub 20180131. doi: 10.1016/j.bone.2018.01.022. PubMed PMID: 29360619.

17. Whyte MP. xHypophosphatasia - aetiology, nosology, pathogenesis, diagnosis and treatment. Nat Rev Endocrinol. 2016;12(4):233–46. Epub 20160219. doi: 10.1038/nrendo.2016.14. PubMed PMID: 26893260.

18. Lundgren T, Westphal O, Bolme P, Modeer T, Noren JG. Retrospective study of children with hypophosphatasia with reference to dental changes. European Journal of Oral Sciences. 1991;99(5):357–64.

19. Reibel A, Maniere MC, Clauss F, Droz D, Alembik Y, Mornet E, Bloch-Zupan A. Orodental phenotype and genotype findings in all subtypes of hypophosphatasia. Orphanet J Rare Dis. 2009;4:6. Epub 20090221. doi: 10.1186/1750-1172-4-6. PubMed PMID: 19232125; PubMed Central PMCID: PMCPMC2654544.

20. Foster BL, Ramnitz MS, Gafni RI, Burke AB, Boyce AM, Lee JS, et al. Rare bone diseases and their dental, oral, and craniofacial manifestations. J Dent Res. 2014;93(7 Suppl):7s–19s. Epub 20140403. doi: 10.1177/0022034514529150. PubMed PMID: 24700690; PubMed Central PMCID: PMCPMC4107543.

21. Kramer K, Chavez MB, Tran AT, Farah F, Tan MH, Kolli TN, et al. Dental defects in the primary dentition associated with hypophosphatasia from biallelic ALPL mutations. Bone. 2021;143:115732. Epub 20201104. doi: 10.1016/j.bone.2020.115732. PubMed PMID: 33160095; PubMed Central PMCID: PMCPMC7769999.

22. Bloch-Zupan A. Hypophosphatasia: diagnosis and clinical signs - a dental surgeon perspective. Int J Paediatr Dent. 2016;26(6):426–38. Epub 20160331. doi: 10.1111/ipd.12232. PubMed PMID: 27030892.

23. Collmann H, Mornet E, Gattenlohner S, Beck C, Girschick H. Neurosurgical aspects of childhood hypophosphatasia. Childs Nerv Syst. 2009;25(2):217–23. Epub 20080904. doi: 10.1007/s00381-008-0708-3. PubMed PMID: 18769927.

24. Durussel J, Liu J, Campbell C, Nam HK, Hatch NE. Bone mineralization-dependent craniosynostosis and craniofacial shape abnormalities in the mouse model of infantile hypophosphatasia. Dev Dyn. 2016;245(2):175–82. Epub 20151228. doi: 10.1002/dvdy.24370. PubMed PMID: 26605996.

25. Houston B, Seawright E, Jefferies D, Hoogland E, Lester D, Whitehead C, Farquharson C. Identification and cloning of a novel phosphatase expressed at high levels in differentiating growth plate chondrocytes. Biochim Biophys Acta. 1999;1448(3):500–6. doi: 10.1016/s0167-4889(98)00153-0. PubMed PMID: 9990301.

26. Houston B, Stewart AJ, Farquharson C. PHOSPHO1-A novel phosphatase specifically expressed at sites of mineralisation in bone and cartilage. Bone. 2004;34(4):629–37. doi: 10.1016/j.bone.2003.12.023. PubMed PMID: 15050893.

27. Roberts S, Narisawa S, Harmey D, Millán JL, Farquharson C. Functional involvement of PHOSPHO1 in matrix vesicle-mediated skeletal mineralization. J Bone Miner Res. 2007;22(4):617–27. doi: 10.1359/jbmr.070108. PubMed PMID: 17227223.

28. Liu X, Hasegawa T, Sakakibara M, Yamamoto T, Haraguchi-Kitakamae M, Ishizu H, et al. Histochemical assessment of the anabolic effects of abaloparatide on the femoral metaphyses of ovariectomized mice. J Oral Biosci. 2025;67(2):100663. Epub 20250418. doi: 10.1016/j.job.2025.100663. PubMed PMID: 40254128.

29. Zweifler LE, Ao M, Yadav M, Kuss P, Narisawa S, Kolli TN, et al. Role of PHOSPHO1 in Periodontal Development and Function. J Dent Res. 2016;95(7):742–51. Epub 20160325. doi: 10.1177/0022034516640246. PubMed PMID: 27016531; PubMed Central PMCID: PMCPMC4914864.

30. Dillon S, Staines KA, Millan JL, Farquharson C. How To Build a Bone: PHOSPHO1, Biomineralization, and Beyond. JBMR Plus. 2019;3(7):e10202. Epub 20190707. doi: 10.1002/jbm4.10202. PubMed PMID: 31372594; PubMed Central PMCID: PMCPMC6659447.

31. Roberts SJ, Stewart AJ, Sadler PJ, Farquharson C. Human PHOSPHO1 exhibits high specific phosphoethanolamine and phosphocholine phosphatase activities. Biochem J. 2004;382(Pt 1):59–65. doi: 10.1042/bj20040511. PubMed PMID: 15175005; PubMed Central PMCID: PMCPMC1133915.

32. Taillandier A, Domingues C, De Cazanove C, Porquet-Bordes V, Monnot S, Kiffer-Moreira T, et al. Molecular diagnosis of hypophosphatasia and differential diagnosis by targeted Next Generation Sequencing. Mol Genet Metab. 2015;116(3):215–20. Epub 20150930. doi: 10.1016/j.ymgme.2015.09.010. PubMed PMID: 26432670; PubMed Central PMCID: PMCPMC5257278.

33. Macrae VE, Davey MG, McTeir L, Narisawa S, Yadav MC, Millan JL, Farquharson C. Inhibition of PHOSPHO1 activity results in impaired skeletal mineralization during limb development of the chick. Bone. 2010;46(4):1146–55. Epub 20100104. doi: 10.1016/j.bone.2009.12.018. PubMed PMID: 20053388; PubMed Central PMCID: PMCPMC2842458.

34. Huesa C, Houston D, Kiffer-Moreira T, Yadav MM, Millan JL, Farquharson C. The Functional co-operativity of Tissue-Nonspecific Alkaline Phosphatase (TNAP) and PHOSPHO1 during initiation of Skeletal Mineralization. Biochem Biophys Rep. 2015;4:196–201. doi: 10.1016/j.bbrep.2015.09.013. PubMed PMID: 26457330; PubMed Central PMCID: PMCPMC4594806.

35. Narisawa S, Fröhlander N, Millán JL. Inactivation of two mouse alkaline phosphatase genes and establishment of a model of infantile hypophosphatasia. Dev Dyn. 1997;208(3):432–46. doi: 10.1002/(sici)1097-0177(199703)208:3<432::Aid-aja13>3.0.Co;2-1. PubMed PMID: 9056646.

36. Fedde KN, Blair L, Silverstein J, Coburn SP, Ryan LM, Weinstein RS, et al. Alkaline phosphatase knock-out mice recapitulate the metabolic and skeletal defects of infantile hypophosphatasia. J Bone Miner Res. 1999;14(12):2015–26. doi: 10.1359/jbmr.1999.14.12.2015. PubMed PMID: 10620060; PubMed Central PMCID: PMCPMC3049802.

37. Liu J, Nam HK, Campbell C, Gasque KC, Millán JL, Hatch NE. Tissue-nonspecific alkaline phosphatase deficiency causes abnormal craniofacial bone development in the Alpl(-/-) mouse model of infantile hypophosphatasia. Bone. 2014;67:81–94. Epub 20140709. doi: 10.1016/j.bone.2014.06.040. PubMed PMID: 25014884; PubMed Central PMCID: PMCPMC4149826.

38. McKee MD, Nakano Y, Masica DL, Gray JJ, Lemire I, Heft R, et al. Enzyme replacement therapy prevents dental defects in a model of hypophosphatasia. J Dent Res. 2011;90(4):470–6. Epub 20110106. doi: 10.1177/0022034510393517. PubMed PMID: 21212313; PubMed Central PMCID: PMCPMC3144124.

39. Foster BL, Kuss P, Yadav MC, Kolli TN, Narisawa S, Lukashova L, et al. Conditional Alpl Ablation Phenocopies Dental Defects of Hypophosphatasia. J Dent Res. 2017;96(1):81–91. Epub 20161001. doi: 10.1177/0022034516663633. PubMed PMID: 27582029; PubMed Central PMCID: PMCPMC5347426.

40. Mohamed FF, Phanrungsuwan A, de Oliveira FA, Millán JL, Foster BL. Dentoalveolar defects and impaired alveolar bone healing in a neural crest directed conditional knockout mouse model of hypophosphatasia. Bone. 2025;198:117538. Epub 20250519. doi: 10.1016/j.bone.2025.117538. PubMed PMID: 40398628; PubMed Central PMCID: PMCPMC12880574.

41. Mohamed FF, Chavez MB, de Oliveira FA, Narisawa S, Farquharson C, Millán JL, Foster BL. Perspective on Dentoalveolar Manifestations Resulting From PHOSPHO1 Loss-of-Function: A Form of Pseudohypophosphatasia? Front Dent Med. 2022;3. Epub 20220203. doi: 10.3389/fdmed.2022.826387. PubMed PMID: 36185572; PubMed Central PMCID: PMCPMC9521815.

42. Yadav MC, Simão AM, Narisawa S, Huesa C, McKee MD, Farquharson C, Millán JL. Loss of skeletal mineralization by the simultaneous ablation of PHOSPHO1 and alkaline phosphatase function: a unified model of the mechanisms of initiation of skeletal calcification. J Bone Miner Res. 2011;26(2):286–97. Epub 20100803. doi: 10.1002/jbmr.195. PubMed PMID: 20684022; PubMed Central PMCID: PMCPMC3179344.

43. Bourne LE, Sharma A, Dillon S, Keen J, Jayash SN, Crump N, et al. TNAP and PHOSPHO1 Function Synergistically to Afford Critical Control Over the Mineralization of the Postnatal Murine Skeleton. Adv Sci (Weinh). 2026:e77671. Epub 20260909. doi: 10.1002/advs.77671. PubMed PMID: 42717495; PubMed Central PMCID: PMCPMC13558897.

44. Bowden SA, Foster BL. Alkaline Phosphatase Replacement Therapy for Hypophosphatasia in Development and Practice. Adv Exp Med Biol. 2019;1148:279–322. doi: 10.1007/978-981-13-7709-9_13. PubMed PMID: 31482504.

45. Lira Dos Santos EJ, Mohamed FF, Kramer K, Foster BL. Dental manifestations of hypophosphatasia: translational and clinical advances. JBMR Plus. 2025;9(2):ziae180. Epub 20250106. doi: 10.1093/jbmrpl/ziae180. PubMed PMID: 39872235; PubMed Central PMCID: PMCPMC11770227.

46. van den Bos T, Handoko G, Niehof A, Ryan LM, Coburn SP, Whyte MP, Beertsen W. Cementum and dentin in hypophosphatasia. J Dent Res. 2005;84(11):1021–5. doi: 10.1177/154405910508401110. PubMed PMID: 16246934.

47. Bloch-Zupan A, Vaysse F. Hypophosphatasia: oral cavity and dental disorders. Arch Pediatr. 2017;24(5S2):5S80–5S4. doi: 10.1016/S0929-693X(18)30020-4. PubMed PMID: 29405938.

48. Ohkura N, Nam HK, Liu F, Hatch N. Cranial Neural Crest Specific Deletion of Alpl (TNAP) via P0-Cre Causes Abnormal Chondrocyte Maturation and Deficient Cranial Base Growth. Int J Mol Sci. 2023;24(20). Epub 20231020. doi: 10.3390/ijms242015401. PubMed PMID: 37895082; PubMed Central PMCID: PMCPMC10607232.

49. Logan M, Martin JF, Nagy A, Lobe C, Olson EN, Tabin CJ. Expression of Cre Recombinase in the developing mouse limb bud driven by a Prxl enhancer. Genesis. 2002;33(2):77–80. doi: 10.1002/gene.10092. PubMed PMID: 12112875.

50. Ono N, Kronenberg HM. Mesenchymal progenitor cells for the osteogenic lineage. Curr Mol Biol Rep. 2015;1(3):95–100. Epub 20150716. doi: 10.1007/s40610-015-0017-z. PubMed PMID: 26526380; PubMed Central PMCID: PMCPMC4624419.

51. Rigueur D, Lyons KM. Whole-mount skeletal staining. Methods Mol Biol. 2014;1130:113–21. doi: 10.1007/978-1-62703-989-5_9. PubMed PMID: 24482169; PubMed Central PMCID: PMCPMC5384832.

52. Goring A, Sharma A, Javaheri B, Smith RC, Kanczler JM, Boyde A, et al. Regulation of the Bone Vascular Network is Sexually Dimorphic. J Bone Miner Res. 2019;34(11):2117–32. Epub 20191009. doi: 10.1002/jbmr.3825. PubMed PMID: 31269275; PubMed Central PMCID: PMCPMC6899569.

53. Chavez MB, Chu EY, Kram V, de Castro LF, Somerman MJ, Foster BL. Guidelines for Micro-Computed Tomography Analysis of Rodent Dentoalveolar Tissues. JBMR Plus. 2021;5(3):e10474. Epub 20210303. doi: 10.1002/jbm4.10474. PubMed PMID: 33778330; PubMed Central PMCID: PMCPMC7990153.

