## Supplementary Information Appendix for "Craniofacial abnormalities from TNAP and PHOSPHO1 deletion highlight their dual role in bone and tooth mineralisation"

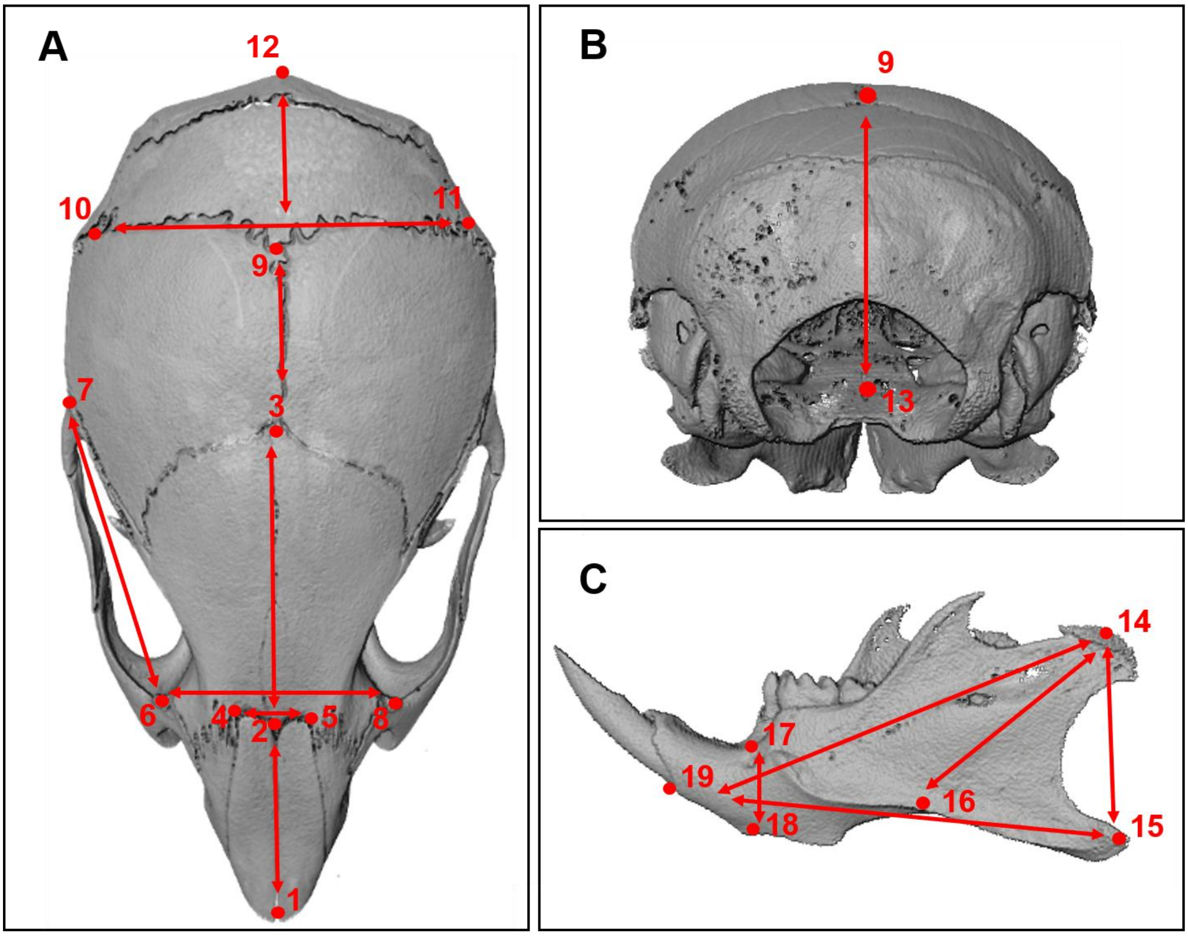


***Supplementary Figure 1. Craniometric measurements from microCT reconstructions.*** *Schematic showing landmarks used to take cranial measurements. (**A**) 1-2: nasal bone length, 1-3: nose length, 2-3: frontal length, 4-5: inner canthal distance, 6-7: zygomatic arch length, 6-8: interorbital distance, 3-9: parietal length, 10-11: skull width, 1-12: skull length. (**B**) 9-13: skull height. (**C**) 14-15: posterior mandible height, 14-16: condylar axis, 14-19: mandible length, 15-19: mandible plain, 17-18: anterior mandible length.*

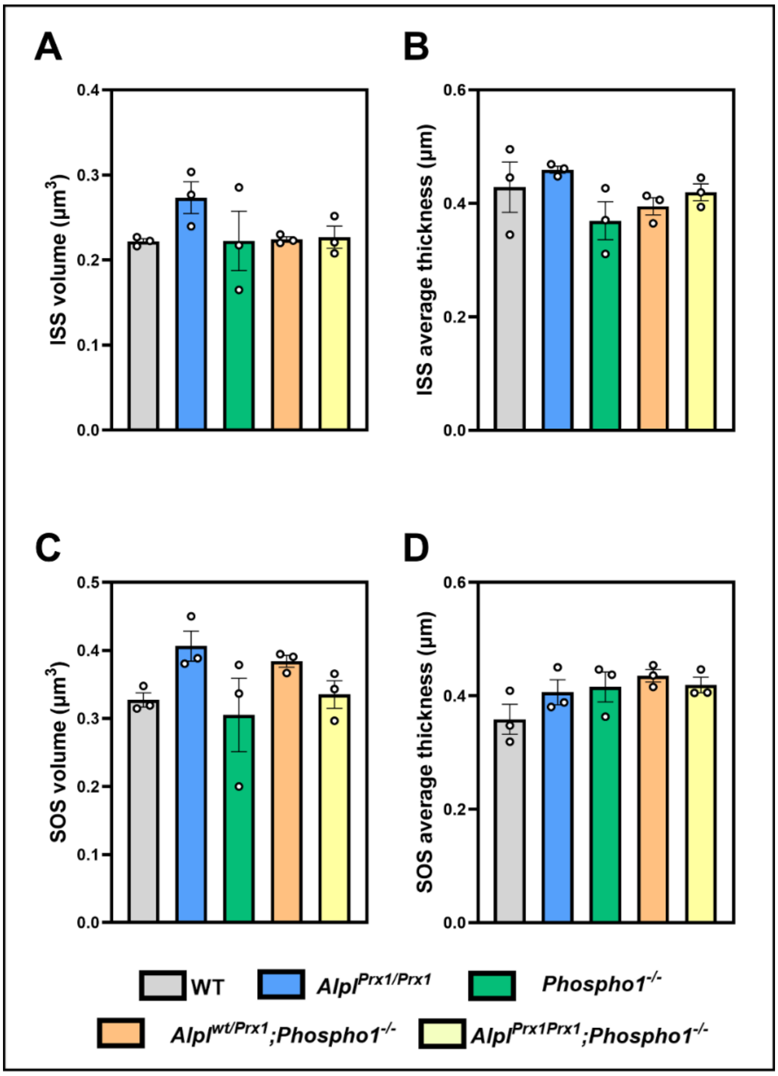


***Supplementary Figure 2. No differences in intersphenoidal synchondrosis (ISS) or spheno-occipital synchondrosis (SOS) formation with Alpl and Phospho1 deletion in 1-day-old mice.*** *ISS (**A**) volume and (**B**) average thickness, and SOS (**C**) volume and (**D**) average thickness in wild-type (WT), Alpl$^{Prx1/Prx1}$, Phospho1$^{-/-}$, Alpl$^{wt/Prx1}$;Phospho1$^{-/-}$ and Alpl$^{Prx1/Prx1}$;Phospho1$^{-/-}$ 1-day-old mice. Data presented as mean ± SEM with points showing individual animals (n=3/genotype).*

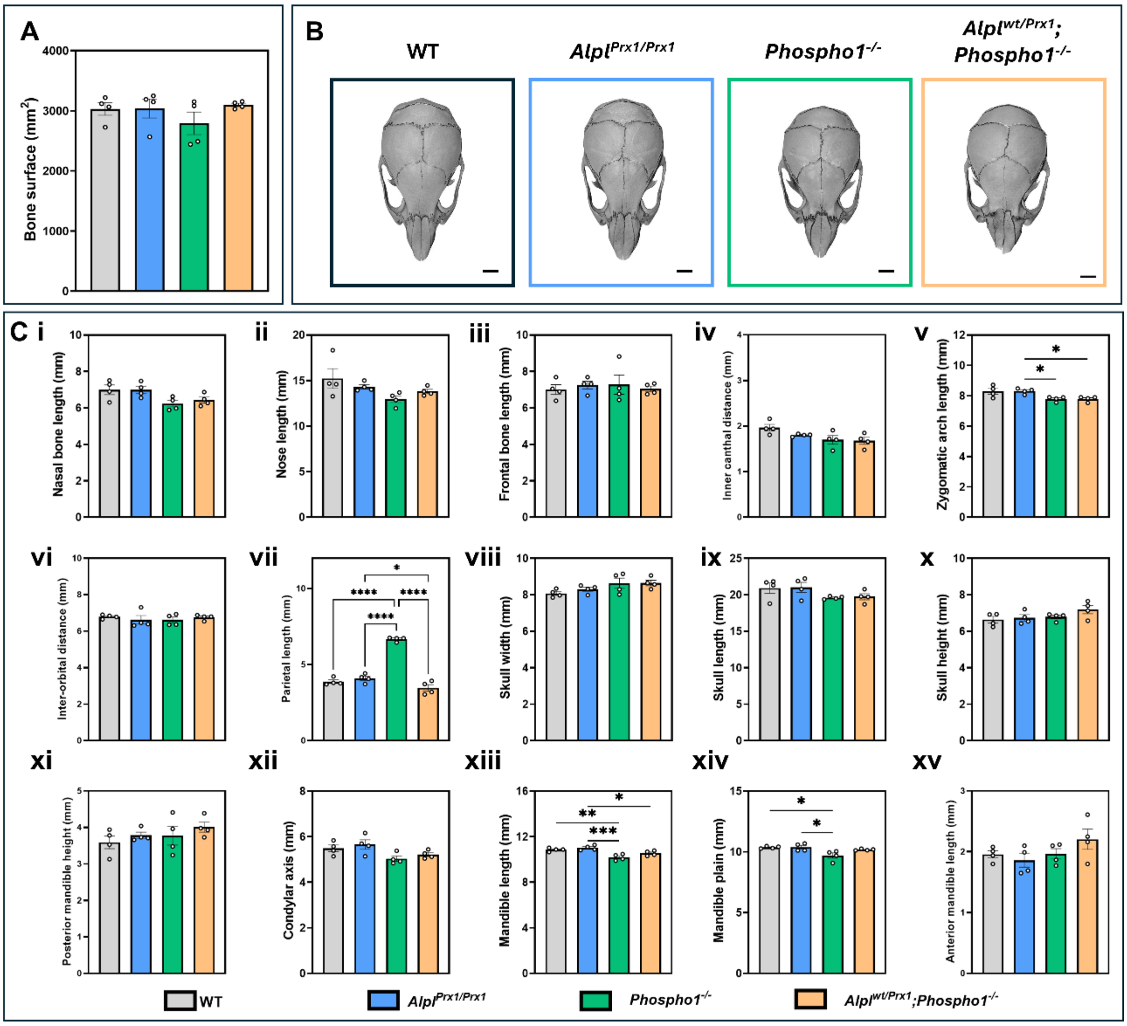


***Supplementary Figure 3. Minimal cranial and mandibular changes in 6-week-old male mice.*** *(**A**) Cranial bone surface area was unchanged. (**B**) Representative 3D-reconstructed images of male skulls of each genotype. (**C**) Anatomical landmark measurements of the skull for (**i**) nasal bone length, (**ii**) nose length, (**iii**) frontal length, (**iv**) inner canthal distance, (**v**) zygomatic arch length, (**vi**) inter-orbital distance, (**vii**) parietal length, (**viii**) skull width, (**ix**) length, (**x**) height. Mandibular landmarks were also measured for (**xi**) posterior mandible height, (**xii**) condylar axis, (**xiii**) mandible length, (**xiv**) mandible plane, (**xv**) and anterior mandible length. Data presented as mean ± SEM with points showing individual animals (n=4/genotype). *= $p<0.05$, **=$p<0.01$, ***=$p<0.001$, ****= $p<0.0001$. Scale bar = 2 mm.*

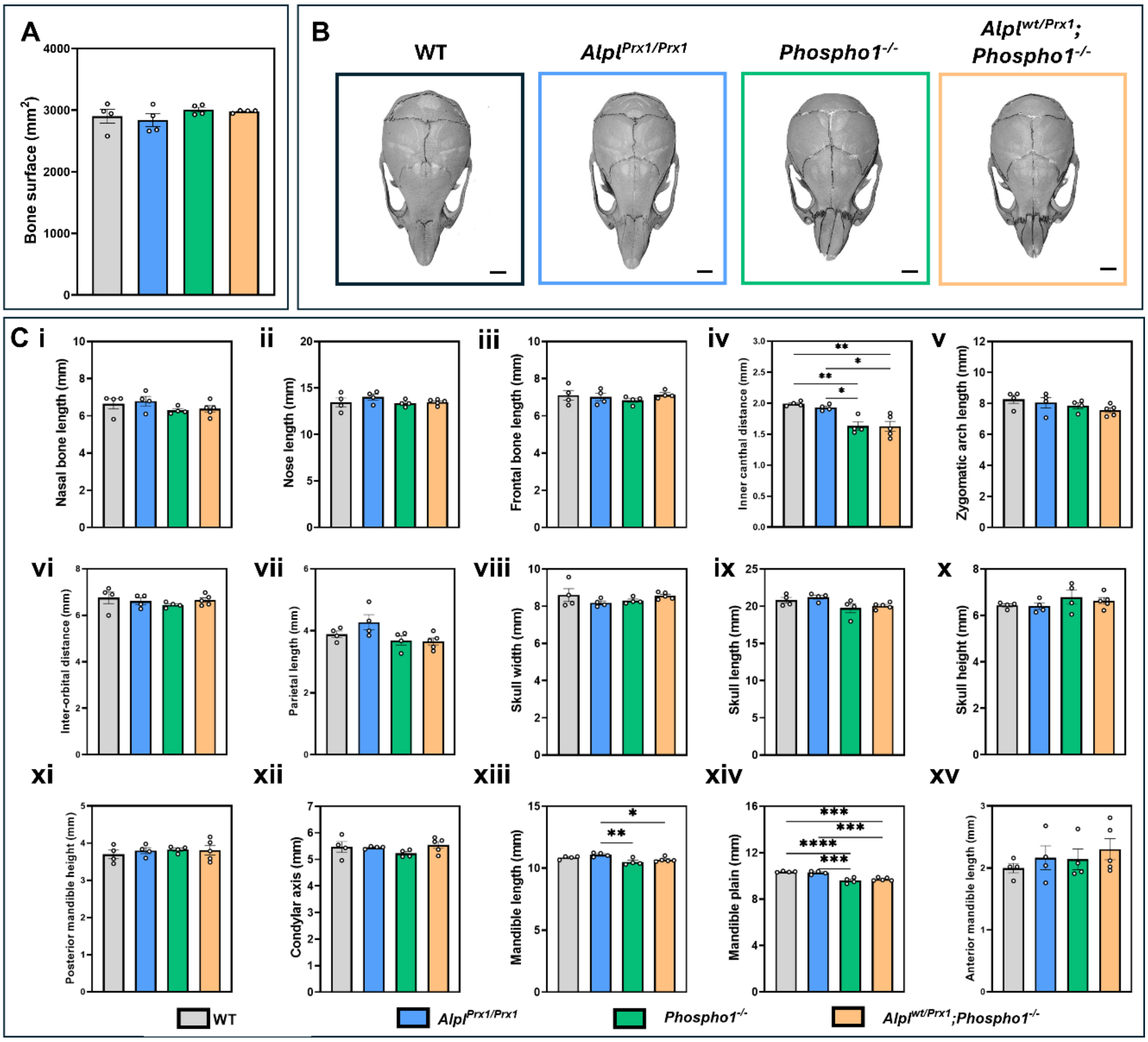


***Supplementary Figure 4. Minimal cranial and mandibular changes in 6-week-old female mice.*** *(**A**) Cranial bone surface area was unchanged. (**B**) Representative 3D-reconstructed images of female skulls of each genotype. (**C**) Anatomical landmark measurements of the skull for (**i**) nasal bone length, (**ii**) nose length, (**iii**) frontal length, (**iv**) inner canthal distance, (**v**) zygomatic arch length, (**vi**) inter-orbital distance, (**vii**) parietal length, (**viii**) skull width, (**ix**) length, (**x**) height. Mandibular landmarks were also measured for (**xi**) posterior mandible height, (**xii**) condylar axis, (**xiii**) mandible length, (**xiv**) mandible plane, (**xv**) and anterior mandible length. Data presented as mean ± SEM with points showing individual animals (n=4-5/genotype). *= p<0.05, **=p<0.01, ***=p<0.001, ****= p<0.0001. Scale bar = 2 mm.*

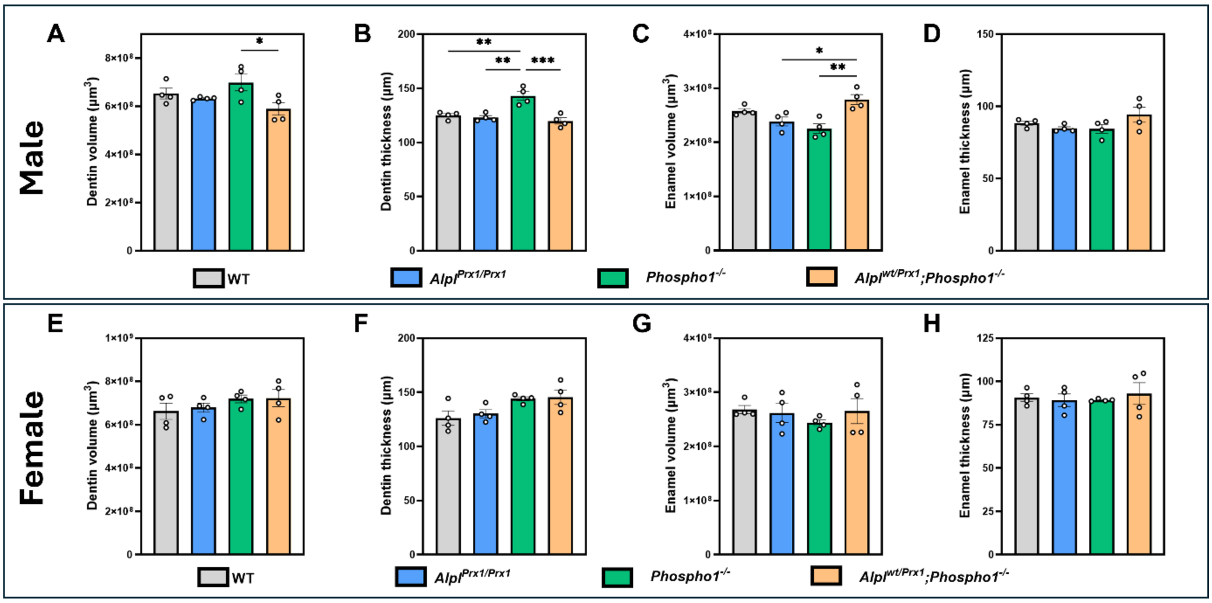


***Supplementary Figure 5. Dentin and enamel changes in 6-week-old mice of both sexes with Phospho1 and Alpl deletion.*** *Micro-computed tomography (µCT) analysis of (**A**) dentin volume (µm$^3$), (**B**) average dentin thickness (µm), (**C**) enamel volume (µm$^3$) and (**D**) average enamel thickness (µm) of the first molar (M1) in male wild-type (WT), Alpl$^{Prx1/Prx1}$, Phospho1$^{-/-}$, and Alpl$^{wt/Prx1}$;Phospho1$^{-/-}$. (**E**) dentin volume (µm$^3$), (**F**) average dentin thickness (µm), (**G**) enamel volume (µm$^3$) and (**H**) average enamel thickness (µm) of the first molar (M1) in female animals. Data presented as mean ± SEM with points showing individual animals (n=4/genotype). *= p<0.05, **=p<0.01, ***=p<0.001, ****= p<0.0001.*
